# Kinetics of multi-substrate in *Beauveriabassiana JUCHE22*

**DOI:** 10.1101/089599

**Authors:** Subhoshmita Mondal, Sarangam Majumdar

## Abstract

The pattern of the biomass specific growth rate is an important focus for many biotechnological applications. Several control strategies have been employed to describe version of the exponential phase of growth of microorganisms. Moreover, in some bioprocesses there is more than one feeding substrates required for the specific growth. Hence, the problem of estimating multiple substrate consumption rates together with the specific growth rate of the microorganism becomes applicable. In this context, the dynamics behaviour of fed-batch process with multiple substrates (dextrose and peptone) and kinetics is further investigated in *Beauveria bassiana*. Different features of the proposed model are assessed by numerical simulation in different scenarios.

## Introduction

*Beauveria bassiana* which has been considered as one of the reliable amongst the group of entomopathogenic fungi is on the focus of investigations in various fields. It is the most favourable and eminent for its anti-pest activity, natural soil borne insect pathogens and are currently being reviewed as an effective alternative to chemical pesticides. Understanding the necessity of finding improved sources of biopesticides for protecting crops as well as environment, the present investigation has been undertaken to study in depth *B. bassiana* starting from its isolation and pattern of growth on the basis of multi-substrate consumption (Mondal et al. 2015). The growth strategy has been optimized using various compatible media. But the consumption of multi-substrate in the media containing carbon source (as Dextrose) and protein or nitrogen source (as Peptone) has gained importance in this investigation.

The expanding biotechnological industry is in need of more efficient, applicable and safe processes to optimize production and improve quality. From a biological view, growth rate is related to substrate consumption rate. Thus, tuning the substrate concentration to achieve growth rate has been the practise of the researchers. The pattern formation describes the mechanism of the biomass concentration in space and time which is an important characteristic feature of estimating multiple substrate consumption rates together with the specific growth rate in microorganism like *Beauveria bassiana.* It shows a large variety of different patterns formed during biomass specific growth rate. The resulting shapes depend on the growth conditions which enhance the complexity of resulting patterns. We have discussed the experiment, mathematical model and the pattern formation from this modelling point of view in the subsequent section.

## Isolation and Culture Conditions

Isolation was done from soil tea plantation areas. The samples were inoculated in SDAY (Sabouraund’s dextrose agar, with 1% of yeast extract) media in petri plates at pH 5.6, 25°C in total darkness for 5 days. About 1gm of each soil sample was inoculated in SDAY (Sabouraund’sdextrose agar, with 1% of yeast extract) media containing dextrose (20 g/l), peptone (10g/l), agar (10g/l) and yeast extract (1%). The best profused fungal inoculums was selected and subcultured in liquid broth with the same SDY media composition to further study the consumption of the substrates at different compositions. From the morphological feature and chemotaxonomic study, it was identified that the spore culture is *Beauveria bassiana* (Published strain, 2015).

## Mathematical Model

We consider biphasic biomass growth. The previously used model to describe dual-substrate fed-batch fermentations accepts a dynamical system in the state-space (Bastin and Dochain 1990; Chang 2003; Dunn et al. 2003).

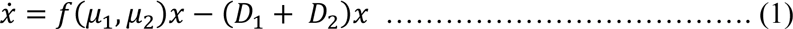

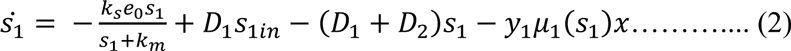

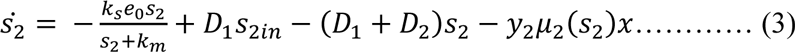

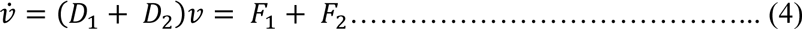

Where,*f*(.) is the specific growth rate, usually the sum or the product of the argument. The state variables are *x* biomass concentration *s_i_* concentration of substrate, and *υ* volume. The specific consumption rates *μ_i_* are unknown nonlinear function of substrates. We consider Haldane kinetics in the consumption rates which have the following form

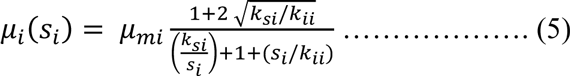

Here, we introduce terms –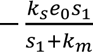 and –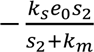 which is consider as perturbation terms in the dynamical system where *k_s_*, *e_0_* and *k_m_* are constant. The parameters *y_i_* are yield coefficients. The other two parameters *s_iin_* are the substrate concentrations in thecorresponding feeding ow. Finally, the *D_i_* dilution ratesare equal to the ratios*F_i_*/*υ*. The substrates may play different roles (Zinn et al. 2004). Both substrates contribute both to growth and production. One substrate is a carbon source mainly affecting growth and the other one a nitrogen source affecting production and product characteristics.

In either case there are mainly two point of views, firstly it is desirable to keep a given specific growth rate *f* = *μ_ref_*, and hence consumption rates *μ_1_* and *μ_2_* corresponding to a desired physiological stateat which the microorganism behaves optimally withrespect to production, does not produce inhibiting products, etc. Secondly, it has been reported in for example (Chang 2003; Kellerhals et al. 1999) that in many instances the ratio 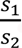 affects the product characteristics, e.g.in PHB production the bioplastic physical properties. Both goals could be achieved by regulating the consumption rates for each substrate using *F*_1,2_. This constitutes the main problem addressed in this work. Note that controller and observer design is subject to the following constraints:

- The only on-line measurable variables are volume and biomass and one of the substrate concentrations.
- The signals are non-negative.
- The yield coefficients *y*_1,2_ and the influent substrate concentrations in are uncertain parameters that, moreover, may vary during the process.
- The specific consumption rates *μ_i_* are not precisely known. We only assume they are Haldane-like nonmonotonous functions, some initial estimation of the maximum consumption rates (an informed guess is enough), estimated upper bounds on their time derivative, and a rough idea of the region (*μ_i_*(*s_i_*),*μ_i_*) where the functions *μ_i_*(*s_i_*) live.

## Result and Discussion

### Experimental study

In first case concentration of dextrose was varied from 14g/l to 25g/l and in second case concentration of peptone was varied from 4g/l to 16g/l. The experimental results showed (Figure 1) the optimum consumption of dextrose provided in the media against the hyphal biomass of 1.1g/l. Concurrently, the optimum consumption of peptone (Figure 2) produced hyphal biomass of 0.5g/l. In case of dextrose, hyphal biomass concentration became constant at 22g/l, whereas, hyphal biomass concentration started to decline after 11g/l peptone concentration.

**Figure 1:**
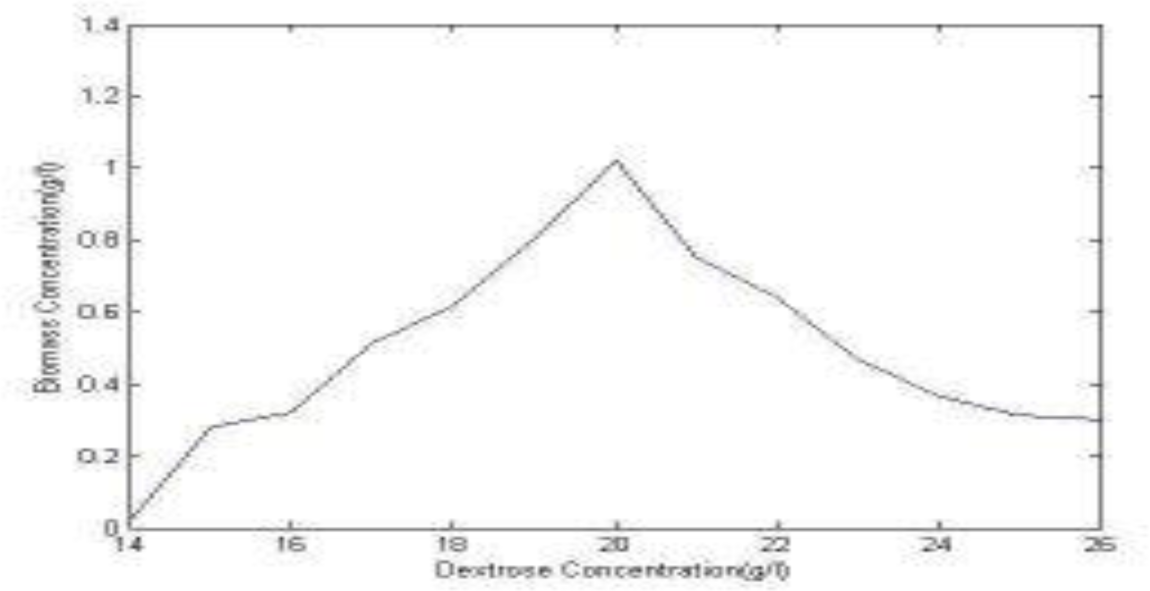
Hyphal biomass concentration (g/l) against different dextrose concentration (g/l).

**Figure 2:**
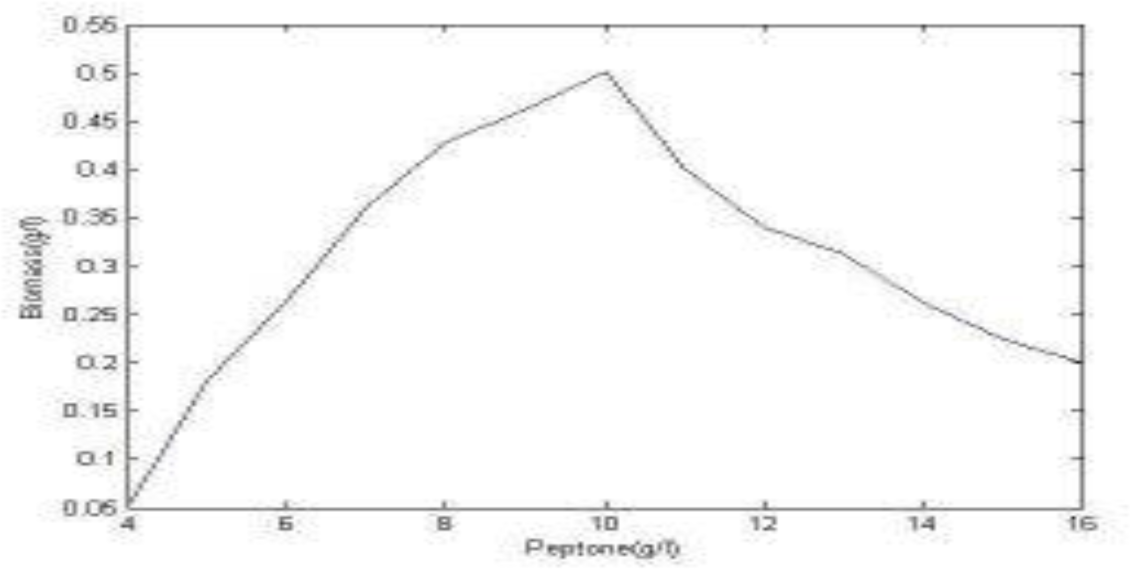
Hyphal biomass concentration (g/l) against different peptone concentration (g/l).

Simple models are discussed (Figure 3) that describe the generation of patterns of the biomass concentration in space and time using numerical technique. The pattern shows the biomass concentration when we are estimating multiple substrate consumption rates together with the specific growth rate of the *Beauveria bassiana* JUCHE22. We get a Gaussian pattern for the biomass concentration which is very much similar to the experimental behaviour. The reliable development of highly complex organisms is an intriguing and fascinating problem. Very complex patterns can be generated in a reproducible way by hierarchical coupling of several such elementary reactions. The complexity of the evolving pattern seems to preclude any mathematical theory. However, by experimental interference with a developing organism it has turned out that the individual steps are fairly independent of each other. The necessity of mathematical models for morphogenesis is evident. Pattern formation is certainly based on the interaction of many components. Since the interactions are expected to be nonlinear, our intuition is insufficient to check whether a particular assumption really accounts for the experimental observation. Models contain often simplifying assumptions and different models may account equally well for a particular observation. This diversity should however be considered as an advantage: multiplicity of models stimulates the design of 1 experimental tests in order to discriminate between the rival theories. In this way, theoretical considerations provide substantial help to the understanding of the mechanisms on which development is based (Berking 1989).

**Figure 3:**
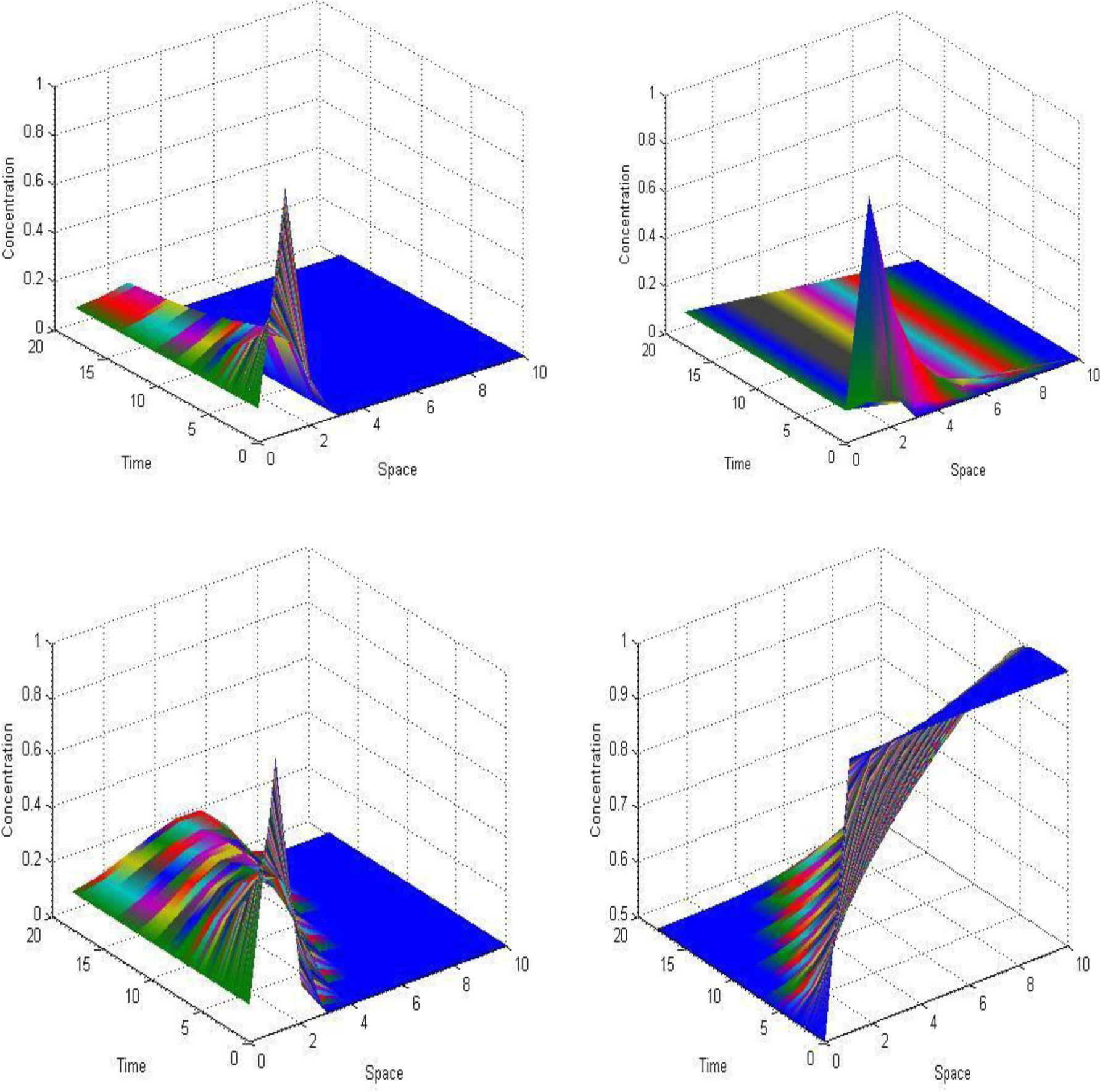
Biomass concentration patterns in space and time.

## Conclusion

The investigational studies of *Beauveria bassiana* JUCHE22 described the optimal consumption rates of the two types of sources provided in the growth media. The dextrose as the carbon source and peptone as the nitrogen source consecutively. The modelled equation considered Haldane kinetics studied the consumption rates along with specific growth rate which suggested there were no inhibiting products during the fed-batch fermentation. Accordingly, biomass concentration patterns were generated following the Gaussian pattern based on the experimental behaviour. This finally explained although there are very high complex patterns, it showed multiplicity stimulating models with the consumption of two substrates in the media.

The authors declare that they have no conflict of interest.

## References

Bastin G, Dochain D (1990) On-line Estimation and Adaptive Control of Bioreactors. Elsevier, Amsterdam.

Berking S (1981) Zur Rolle von Modellen in der Entwicklungsbiologie. In Sitzugsberichte der Heidelberger Akademie der Wissenschaften. Springer-Verlag, Berlin.

Chang DM (2003). The snowball effect in fed-batchbioreactions. Biotechnology Progress 19(3), 1064-1070.doi:10.1021/bp025792a.

Dunn I, Heinzle E, Ingham J et al (2003) Biological Reaction Engineering. Dynamic Modelling Fundamentals with Simulation Examples. Wiley-VCHVerlag.

Kellerhals M.B, Kessler B, Witholt B (1999) Closed-loop control of bacterial high‐cell-density fed batch cultures: Production of mcl-phas by pseudomonasputida kt2442 under single-substrate and cofeeding conditions.Biotechnology and bioengineering 65(3), 306-315.

Mondal S, Datta S, Mukherjee A et al (2015) Studies on isolation, optimization and bioprocess engineering behaviour of entomopathogenic fungi, Beauveria bassiana. Indian Chemical Engineer. doi:10.1080/00194506.2015.1075439 Published strain reference (http://www.ncbi.nlm.nih.gov/nuccore/KT359346.

Xu J, Guo B, Zhang Z et al (2005) A mathematical model for regulating monomer composition of the microbially synthesized polyhydroxyalkanoate copolymers. Biotechnology and bioengineering 90(7), 821-829.

Zinn M, Witholt B, Egli T (2004) Dual nutrient limited growth: models, experimental observations, and applications. Journal of biotechnology 113(1), 263-279.

